# Temporal Synchronization Analysis: A Model-Free Method for Detecting Robust and Nonlinear Brain Activation in fMRI Data

**DOI:** 10.1101/2025.04.21.649810

**Authors:** Suruchi Fialoke, Aniruddha Deb, Kushagra Rode, Vaibhav Tripathi, Rahul Garg

## Abstract

The sluggishness of the fMRI blood oxygenation level dependent (BOLD) signal has motivated the use of block or trial-based experimental designs that rely on the assumption of linearity, typically modeled using the General Linear Model (GLM). But many non-sensory brain regions and subcortical areas do not correspond to such linearities. We introduce a model-free estimation method called Temporal Synchronization Analysis (TSA) which detects significant brain activations across trials and subjects at an individual time point. We validate it across multiple cognitive tasks (combined n=1600). In constrained task stimuli like visual checkerboard paradigms, we discovered novel nonlinearities not reported previously. In model-free task paradigms like listening to naturalistic auditory stimuli, TSA can detect unique stimuli linked quasi-temporal activations across default mode and language networks. Our user-friendly Python toolkit enables cognitive neuroscience researchers to identify stable and robust brain activation across various cognitive paradigms that are challenging to model with current methods.

---

Early cognitive neuroscience studies using fMRI in humans often employed visual tasks because these tasks were well-studied in primates and rodents, providing a reliable validation against established findings in the field. Blood Oxygenated Level Dependent (BOLD) responses in the visual cortex were mostly linear (Boynton et al., 1996; Dale & Buckner, 1997), although even early studies reported nonlinearities with certain stimulations <u>(Friston et al., 1998; Greicius & Menon, 2004)</u>. Since fMRI does not capture a direct neuronal signal <u>(Logothetis, 2008)</u>, the estimation of Hemodynamic Response Function (HRF) to infer neuronal responses from BOLD signals was required. As responses in the visual cortex mostly presented a linear time-invariant (LTI) structure; the General Linear Method (GLM) technique was used under the LTI assumption for their investigation. The GLM analysis worked very well for early sensory and motor regions and was widely adopted in standard analysis packages like FSL <u>(Jenkinson et al., 2012)</u>, SPM (Friston et al., 1994), and AFNI <u>(Cox, 1996)</u>. Although each package allows for the creation of individual-specific HRFs, and some groups ran preliminary tasks for the computation of such HRFs, the double gamma HRF became the standard in the field. As investigators expand their research into higher-order regions across the brain by investigating increasingly complex task paradigms, the LTI assumption often paints an incomplete picture. Studies have shown the poor test-retest reliability of fMRI studies <u>(Elliott et al., 2020)</u> which could potentially be attributed to various factors including scanner differences, BOLD sequence differences, but also to individual-level variability, which such an LTI assumption does not take into consideration <u>(Monti, 2011; Rio et al., 2013).</u> Many studies have reported nonlinearities <u>(Aguirre et al., 1998;</u> <u>Bandettini et al., 2002; Handwerker et al., 2004)</u>, lagged responses, and unconventional peak responses <u>(Bailes et al., 2023; Lewis et al., 2018)</u>.

With the improved computational speeds and easy availability and sharing of statistical methods along with the sharing of large-scale fMRI datasets like the Human Connectome Project (HCP), Nathan Kline Institute Rockland Sample (NKIRS) dataset, etc., we now have the ability to go beyond such a constrained model like GLM. In this paper, we outline a novel model-free, non-parametric method, called Temporal Synchronization Analysis (TSA) <u>(Tripathi & Garg, 2022)</u> and present a user-friendly Python toolbox that facilitates the application of this approach to existing fMRI datasets. The benefit of the TSA method over GLM is that it does not require the assumption of an HRF, is inherently model-free, and instead of a single “Active/not-active” value, it gives “a time course of significant BOLD activity” in response to a stimulus. It also allows researchers to discern subtle differences in the time course between different presentations of the same stimuli (non-LTI behavior), different experiment conditions, different groups of subjects, and a combination of these. Since there is no assumption of the HRF, it can be applied across the brain and subcortical systems and can allow cognitive neuroscientists to find interesting patterns of activity otherwise missed using a constrained model.

**Figure 1.**
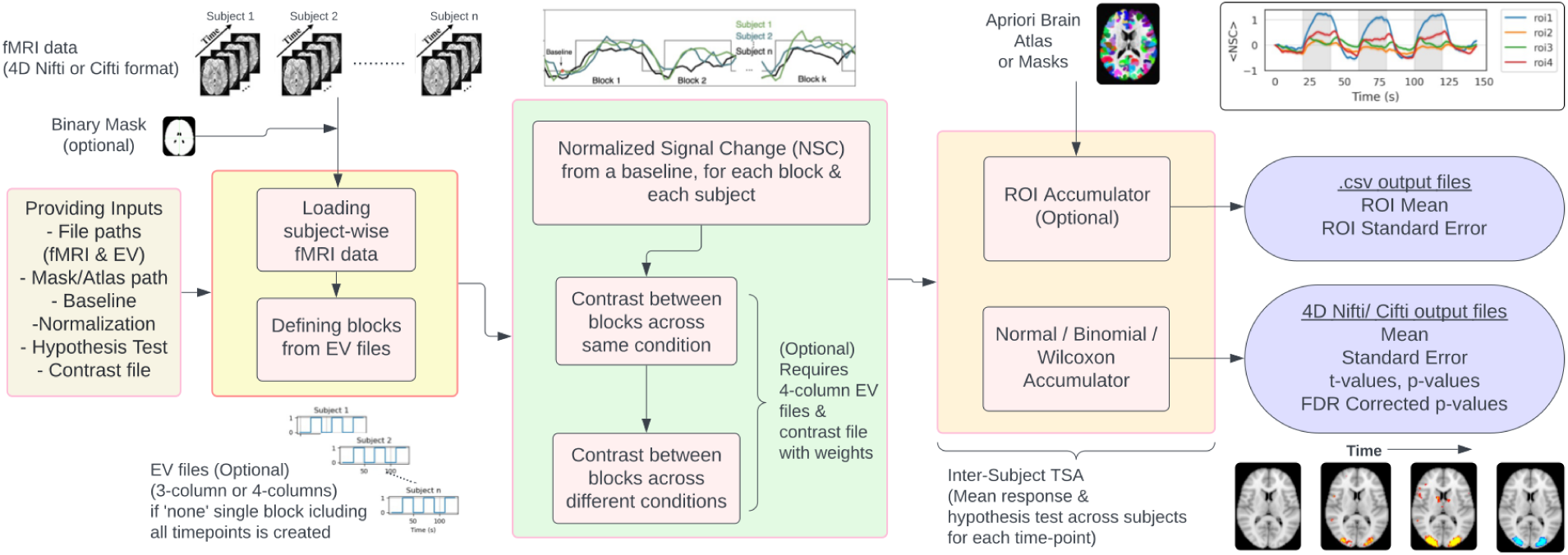
TSA Workflow: An illustrative overview of a Python-based toolkit for Temporal Synchronization Analysis (TSA) across Subjects from fMRI BOLD signal: from subject-wise data loading and block definition to signal accumulation and statistical testing, culminating in comprehensive output files for interpretation.

The key approach of the current method is that different subjects act as independent and identically distributed random variables (IID) and statistical analysis is performed at the time of the BOLD response after stimulus presentation. The model allows for flexible use of different statistical distributions (normal, nonparametric, binomial, and Wilcoxon) to test for significance at each time point. Although given the method, it would be preferable to present stimulus to the subjects in an identical manner, it is not required. The toolkit allows for the reanalysis of existing datasets to get new findings and develop new paradigms to investigate more complex behaviors in naturalistic conditions <u>(Sonkusare et al., 2019)</u>.

In this paper, we highlight the method and the protocol and apply it to large-scale datasets across various cognitive paradigms like investigation of nonlinear brain responses to a simple visual checkerboard task from the NKIRS (n= 317), working memory task from the HCP (n=1200), and complex naturalistic audiovisual film called Pieman from the Narratives dataset (n=82). We report various interesting patterns of activation missed using standard GLM approaches to demonstrate the utility of the TSA approach to investigate data in a model-free manner at an individual time unit level.

## Results

### Visual Checkerboards Task Results

The results from the TSA of checkerboard viewing by 317 subjects are shown in Figure 2 and Figure 3. Figure 2A displays the brain activation maps for FDR-corrected p-values along with three defined regions of interest (ROIs) marked in the brain across several time points. These FDR-corrected p-values are generated as standard output by the TSA Python toolkit. It can be observed from the brain maps that the activation in the three ROIs (Visual Cortex region) increases when the stimulus is presented to the participants. The fixation blocks then resulted in a gradual decrease in the activation in these regions across participants. This accurately depicts the involvement of the three ROIs in the visual processing task involved in the NKI-checkerboard data.

**Figure 2:**
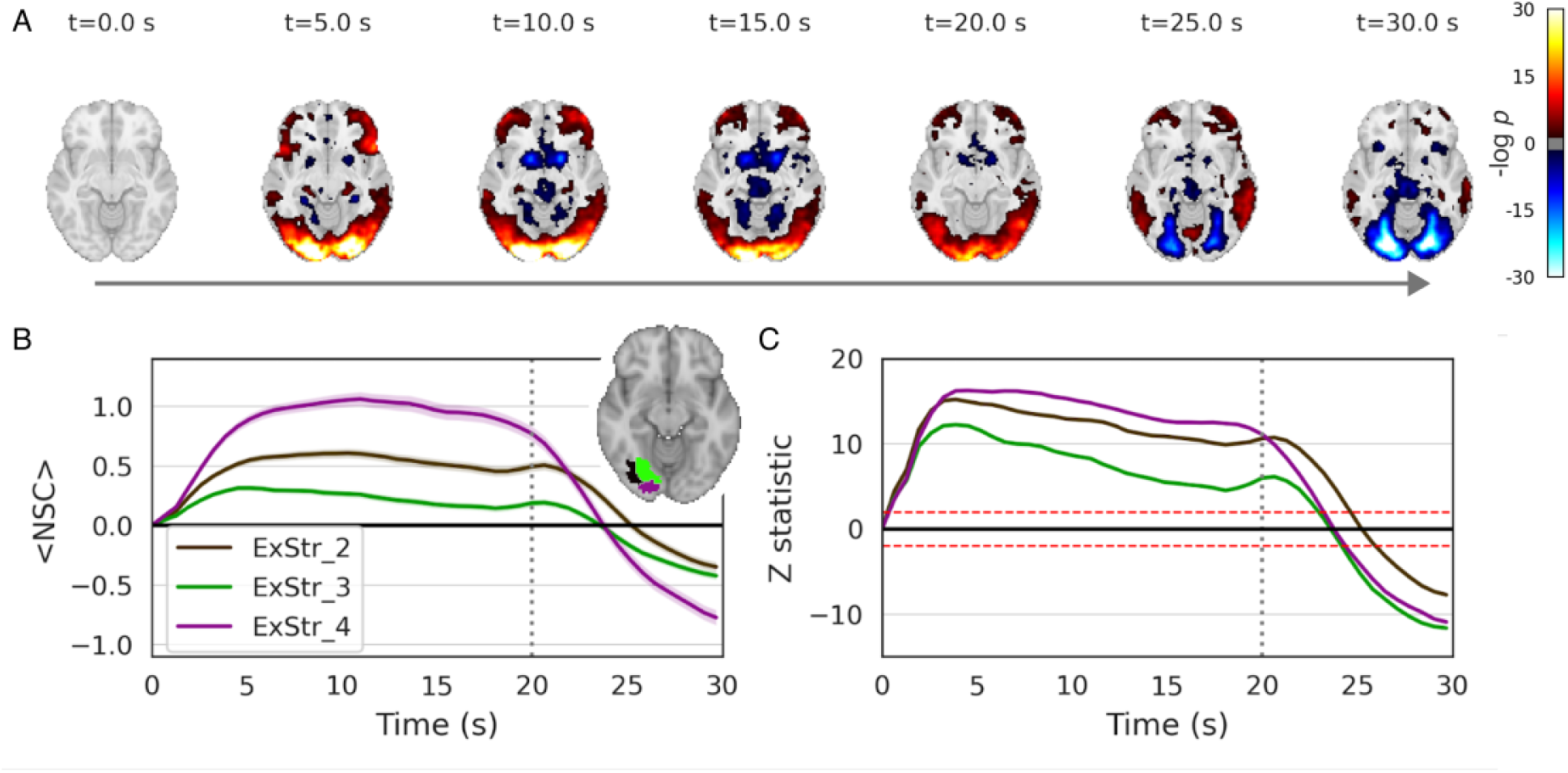
TSA of responses to a visual checkerboard task, across 317 adults from the NKI dataset. (**A)** False discovery rate (FDR)-corrected p-values on a logarithmic scale at specified time intervals, reveals significant synchronized BOLD response in the visual cortex. (**B)** Time-course graph of the mean Normalized Signal Change (NSC) in three defined regions of interest (ROIs) within the visual cortex according to the Schaffer Cortical Atlas. Shaded regions indicate the variability with standard error. Inset: Visualization of the three targeted visual cortex ROIs superimposed on the standardized MNI brain template, providing a spatial reference for parts B and C. (**C**) Time-course representation of z-statistics for the aforementioned ROIs, offering a standardized measure of deviation from expected brain activity. The time course changes are non-homogeneous across the various extrastriate regions during the checkerboard block followed by a small increase during the fixation onset for extrastriate regions 3 and 4. We also observed changes in various prefrontal regions and subcortical structures during the checkerboard condition.

**Figure 3:**
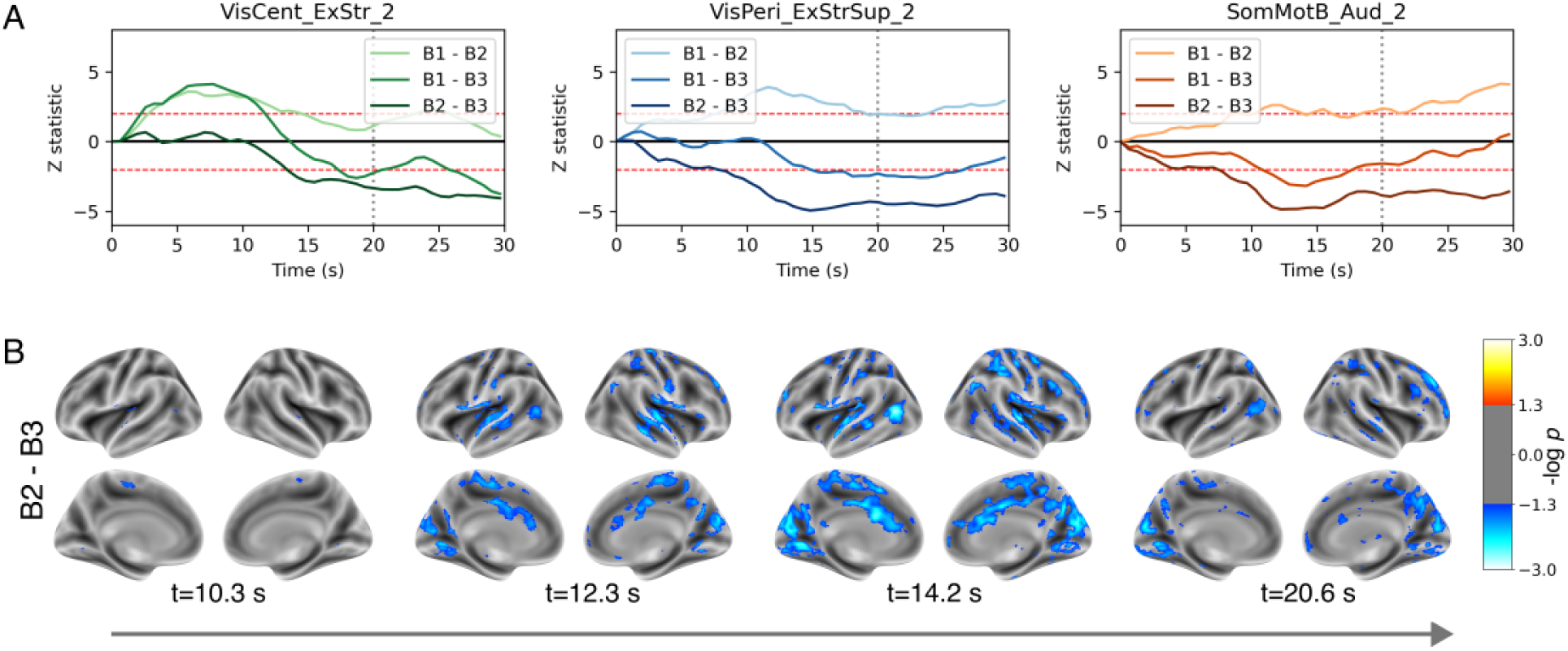
TSA results showcasing differences between blocks of the visual checkerboard task across 317 participants. (**A)** ROI analysis of block contrasts “Block 1 - Block 2 (B1-B2)”, “Block 1 - Block 3 (B1-B3)”, and “Block 2 - Block 3 (B2-B3)” in three ROIs from the Schaefer atlas, showing z-statistics over time. The first ROI is from the primary visual network, whereas the second and third are ROIs with the strongest response for B2-B3 contrast. (**B)** FDR-corrected p-values for the contrast “Block 2 - Block 3 (B2-B3)” for select time points are shown. Although visual checkerboard is assumed to be an LTI experiment with similar results across trials, we show here that this is not the case. Comparing across blocks we see that the first block showed the strongest activations which reduces across the subsequent blocks.

Figures 2B and 2C are time series graphs depicting the mean NSC and the Z statistic, respectively. The ‘roi_mean’ and ‘roi_stderr’ files generated by the toolkit are used to generate the plots. We see a large degree of convergence between the results observed using the TSA and GLM approach (Figures S1-S3). We also observed changes in the prefrontal and subcortical structures during the checkerboard block. The activations at a single TR resolution allow us to see the minor heterogeneities in a paradigm established as a visual checkerboard. We observe progressive decreases in the extrastriate 3 and 4 areas followed by a jump in activation during the block offset and a subsequent decline during the fixation block. Figures 3, S2, and S3 highlight the differences in activations across various blocks with the first being the strongest activator with a reduction in activity during the subsequent blocks. As highlighted in Figure S4, we also see intriguing dynamics of transient changes across the salience network and the somatomotor network where we see an initial deactivation during the first 10 sec followed by a return to baseline.

### TSA contrast between Working Memory Tasks

We then analyzed the TSA contrast between working memory tasks from the HCP dataset. Here, the participants are performing a 2-back and a 0-back working memory paradigm. Figures 4A and S5 show the FDR-corrected p-values for contrasts at selected time points, highlighting the statistically significant differences in brain activation between the 2-back and 0-back conditions over time. These results indicate specific moments where the working memory load (2-back) elicited significantly different neural responses compared to the baseline condition (0-back). The ROI analysis shows z-statistics (Figure 4B) and NSC values (Figure S6A) over time for the 2-back vs. 0-back contrasts in the three ROIs with the lowest z-statistics from the Schaefer atlas. Notably, all these ROIs fall within the Visual Network, despite not being pre-selected based on their network affiliation. This finding suggests that regions within the visual network show significantly lower visual cortex activation in the 2-back memory task compared to the 0-back task. In contrast, the ROIs with the highest z-statistics (Figure 4C) from the Schaefer atlas all fall within the cognitive control network which has been the canonical area for the maintenance, encoding, and retrieval of working memory. We highlight that the TSA method allows to compute contrast between task conditions just like GLM and can be used to extract novel insights into how information maintenance in working memory probably results in reduced visual activations at the onset of the block which slowly comes back to baseline.

**Figure 4:**
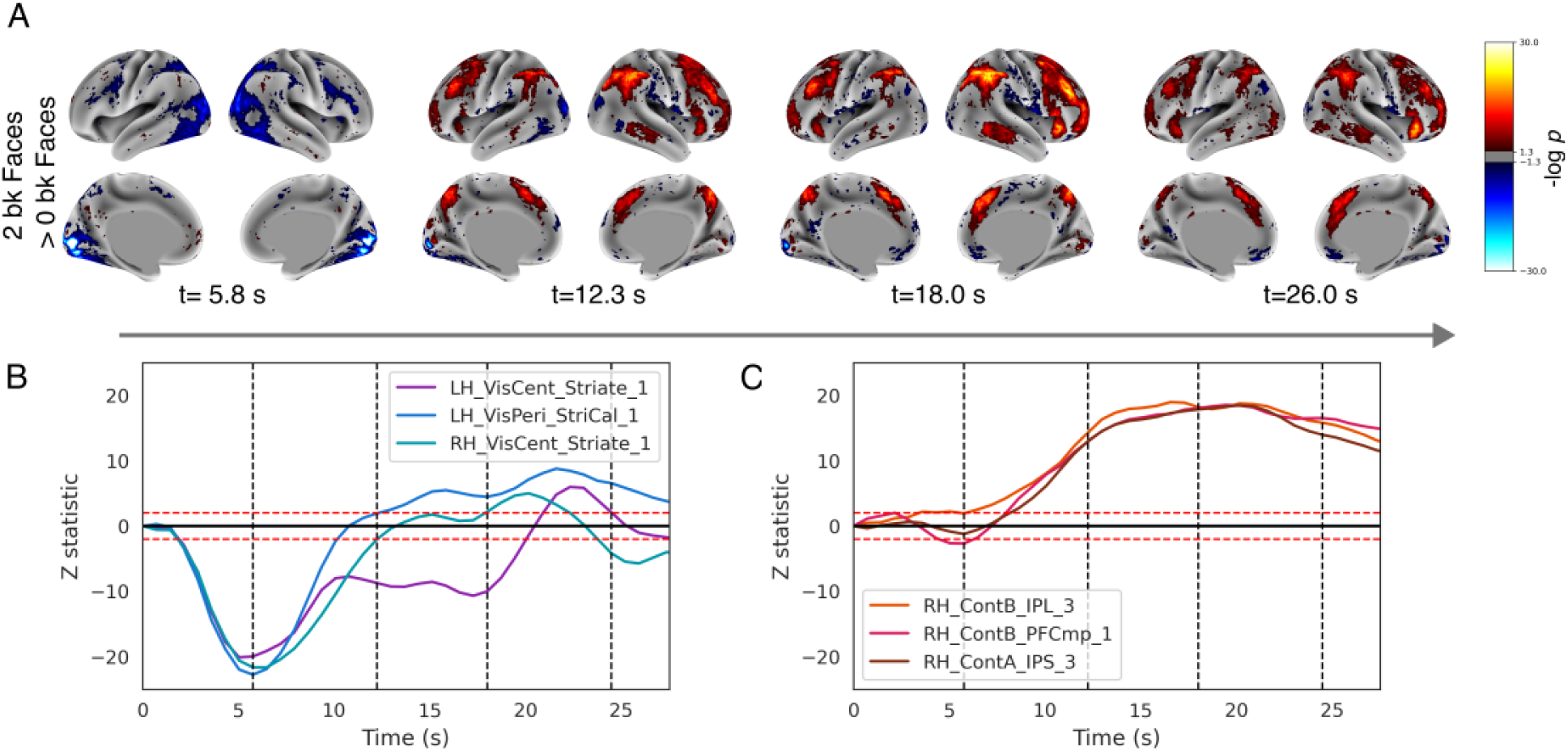
TSA results showcasing differences between working memory task (faces visual stimuli) contrast of 2 back versus 0 back. (**A)** FDR-corrected p-values for the contrast for select time points are shown. (**B)** ROI analysis showing z-statistics over time for the 2-back vs. 0-back contrasts in three ROIs with the lowest z-statistics from the Schaefer atlas, all incidentally within the Visual Network. **C.** ROI analysis shows z-statistics in three ROIs with the strongest 2-back vs. 0-back contrasts over time, remarkably all appearing within the Cognitive Control Network (Sheaffer Atlas).

### TSA of a story listening paradigm from Pieman Narrative Dataset

We wanted to analyze how TSA would be applicable to a naturalistic stimuli paradigm. We analyzed the activations using TSA across participants who listened to the Pieman story (n=82) and found interesting dynamics of activity across the story listening paradigm. We saw consistent changes in activations across participants and dynamical activity patterns across regions in the language network, prefrontal and temporal cortex (Figure 5). Different parts of the story elicited strong synchronization of activity in the frontal operculum, and prefrontal cortex which was distinct from the activity in the temporal cortex (contrasting the time series in Figure 5A and 5B).

**Figure 5:**
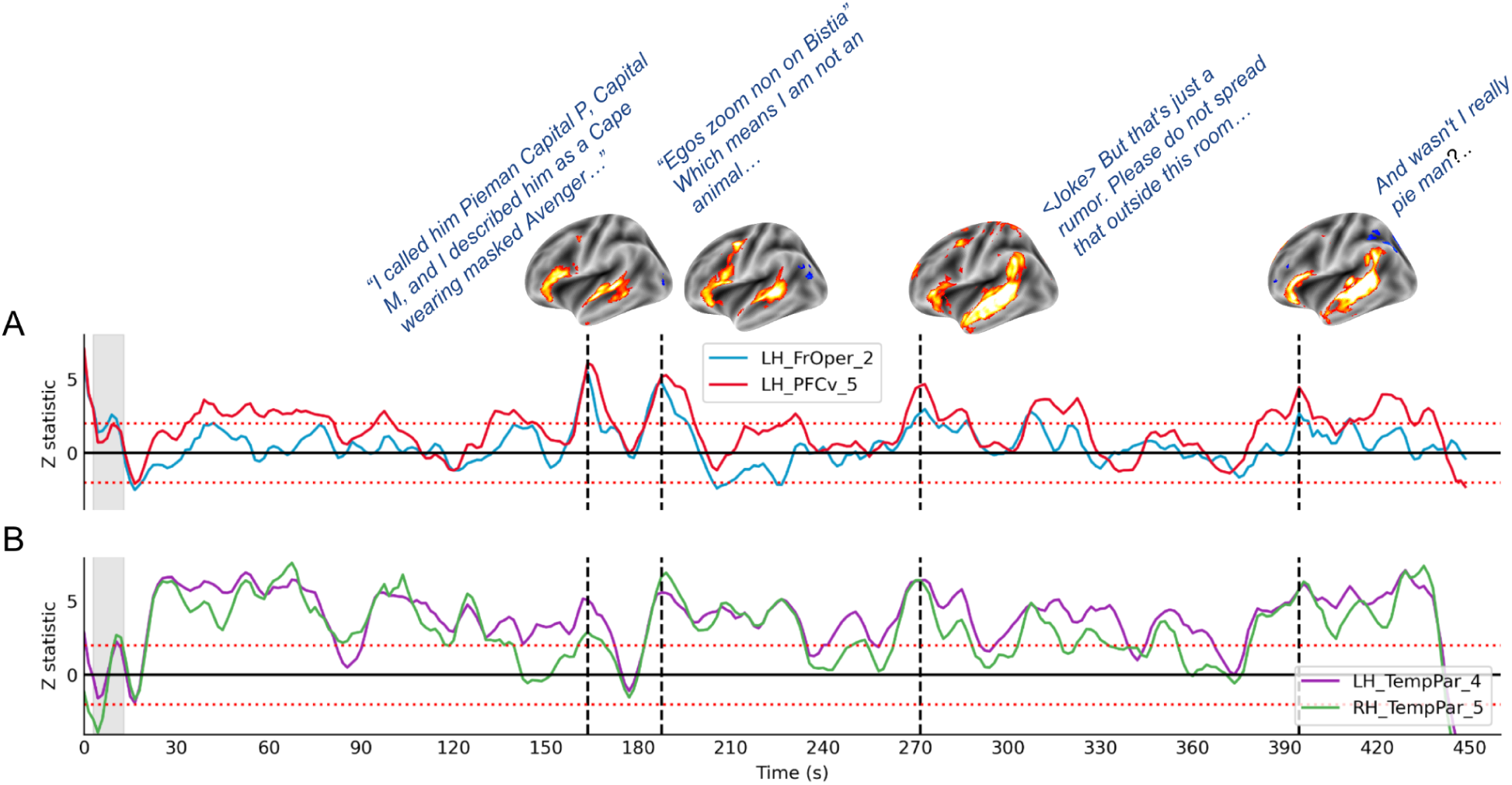
TSA results for the Pieman story listening task across 82 participants. FDR-corrected p-values are displayed for selected time points, along with preceding subtitles. TSA activation (z-statistic) is shown across language processing ROIs, including (B) Broca’s area in the left hemisphere and (C) Wernicke’s area in the left and right hemispheres. A supplementary movie containing all FDR-corrected p-value volumes, along with subtitles and audio, is provided in the Supporting Information (SI).

The activity in default network regions (precuneus, medial PFC, and inferior parietal lobule) varied significantly across the parts of the story (see time points at 184.5 sec and 220.5 sec in Figure 6A). Different parts of the story elicited changes in activity across large-scale brain networks. Even regions within the same network showed heterogeneity in activations (Figure 6B and 6D) suggesting significant potential for discovery in such datasets using a method like TSA.

**Figure 6:**
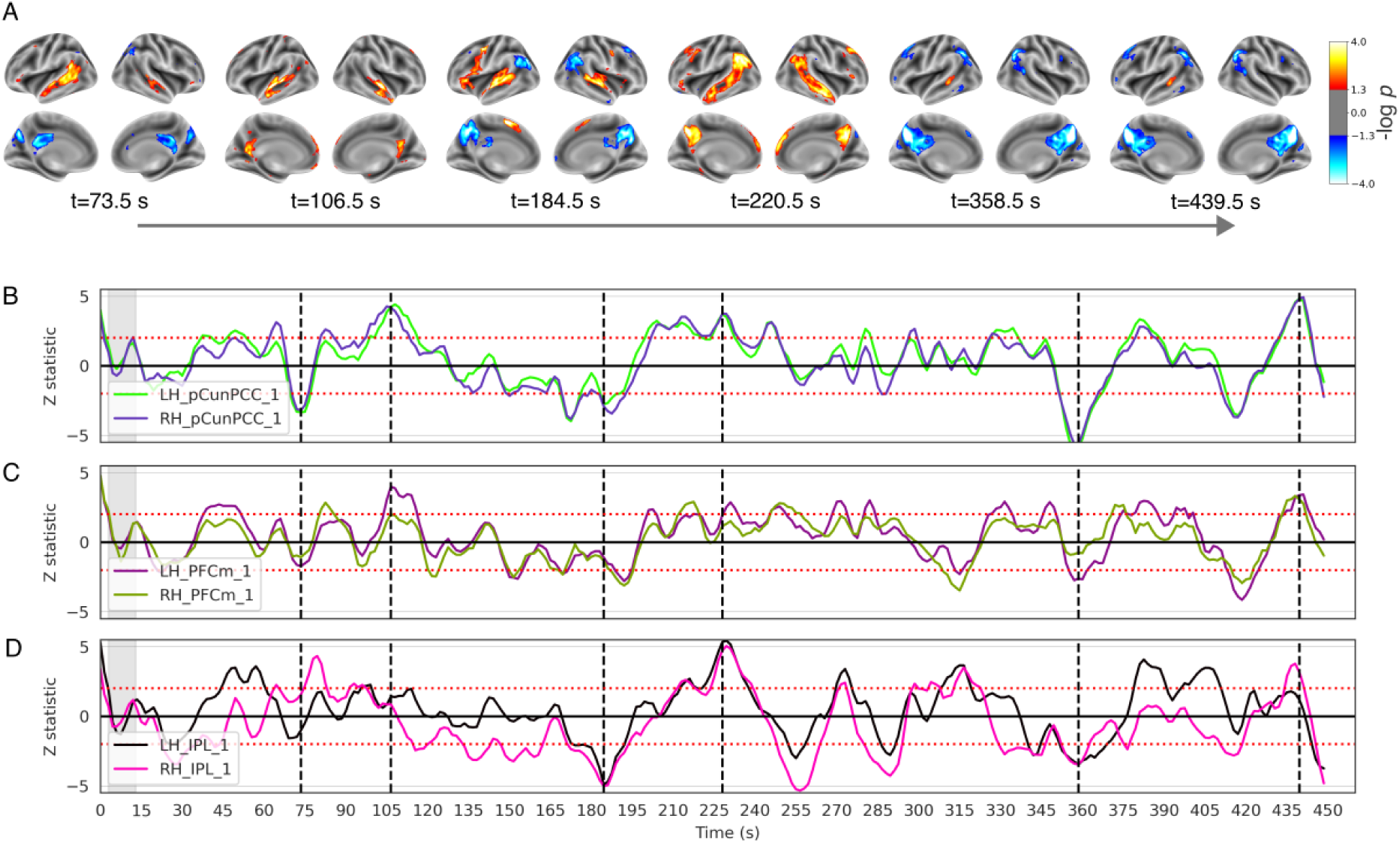
TSA results of Pieman story listening task across 82 participants. (**A)** FDR-corrected p-values for select time points are shown. (**B, C, D)** TSA activations (z-statistic) across various ROIs in the default network (precuneus, medial prefrontal cortex, inferior parietal lobule) for both hemispheres during the story listening task. We see slight variations in activation profiles across the various default network regions especially between the inferior parietal lobule and precuneus/medial PFC.

## Discussion

In this study, we provide a toolkit for a newly developed method called Temporal Synchronization Analysis to analyze fMRI data without the assumptions of hemodynamics which can allow for a finer understanding of different spatiotemporal patterns of activations to a given stimuli across individuals not possible in earlier model constrained methods. We validate the method with three large-scale datasets across block-based tasks (visual checkerboard and working memory) and unconstrained naturalistic stimuli datasets. We discuss the various findings here and the ramifications of using such a tool for better understanding and mapping of brain function using fMRI-based data.

Studies have reported nonlinearities across BOLD signals throughout the brain from the early years of fMRI analysis but it was hard to know whether it was neuronal or hemodynamic <u>(Bandettini et al., 2002; Birn et al., 2001; Friston et al., 1998, 2000;</u> <u>Vazquez & Noll, 1998; Wager et al., 2005)</u>. Various models like the Volterra, Balloon-based, and event-based protocols have tried to model nonlinearities with limited success. Even in recent years, researchers have challenged the notions of LTI assumptions in BOLD data <u>(Chen et al., 2020, 2023)</u> with the presence of hemodynamic response function across various cortical and subcortical regions that departed significantly from the double gamma standard HRF <u>(Lewis et al., 2016, 2018)</u>. Although the evidence is substantial in favor of incorporating nonlinearities in neuroimaging analysis pipelines, existing analysis tools like FSL, and SPM require substantial effort to incorporate them. Our TSA method makes it easier to run a model-free variation which allows for interesting findings across spatiotemporal patterns of activations for a range of cognitive tasks, three of which we explored - visual checkerboard, working memory, and story listening.

For the visual checkerboard condition in the NKIRS dataset, we observed the expected properties in the visual system as many studies have already shown <u>(Boynton et al.,</u> <u>1996</u>). TSA also supplements this information with time series resolution which highlights some interesting dynamics, first, the higher-order extrastriate regions slowly decrease in the BOLD activation during the checkerboard block followed by a sudden increase during the fixation onset followed by a gradual decrease which other studies have also highlighted <u>(de Zwart et al., 2009)</u>. Second, we also found activations in the prefrontal cortex and deactivations in the subcortical areas during the checkerboard stimulation. The activations, albeit weak, become significant when observed consistently across the large number of subjects utilized in the dataset. We also observed non-linearities of activations across multiple presentations for the checkerboard condition, again highlighting the limitations of the GLM model which the TSA approach can overcome.

Then, we analyzed a higher-order working memory task which has been shown to activate regions in the prefrontal cortex and the frontoparietal control network <u>(Assem et al., 2020; Somers et al., 2021)</u>. We demonstrated that the TSA can be utilized to compute contrast across conditions just like GLM but also gives out the temporal structure of the differences across these conditions. This enables us to find that during the working memory block onset (2-back) the visual cortical activity reduces which comes back up after 8-10 secs probably highlighting the role of cognitively demanding tasks in reducing BOLD activity in other areas of the brain. Though studies have shown the role of sustained attention in the reduction of visual responses <u>(Ling & Carrasco, 2006)</u>, we were able to demonstrate it for a working memory paradigm.

The tool also presents a compelling use case for the analysis of the naturalistic stimuli dataset. The field of neuroscience has been lately gravitating towards the use of ecologically valid conditions and naturalistic stimuli like movie watching and audio listening <u>(Sonkusare et al., 2019)</u>. We analyzed the activations during a story listening paradigm and were able to extract interesting dynamics centered in the language network and the default network. We were able to estimate activations across parts of the language network in the temporal and the prefrontal cortex (frontal operculum and dorsolateral prefrontal cortex) with discernible differences in activations across the various regions for specific parts of the story. We also observed differences in activations in subregions of the intrinsic processing default network - the inferior parietal lobule usually associated with the thinking about others or Theory of Mind (DiNicola et al., 2020; Saxe & Kanwisher, 2003) and precuneus and medial PFC associated with episodic projections or scene construction (Buckner & DiNicola, 2019; Du et al., 2024; Yeo et al., 2011). In recent years there has been a whole slew of newer analysis methods to analyze ecologically valid and naturalistic stimuli experiments like the inter-subject correlation (ISC) <u>(Hasson, 2004; Hasson et al., 2008)</u>, inter-subject functional correlation (ISFC) <u>(Simony et al., 2016)</u>, and sequence learning models <u>(Baldassano et al., 2017)</u> which expanded the repertoire of methods that are not dependent on the properties of the HRF and limited by model assumptions. TSA adds to this significantly allowing a single TR resolution of the activity happening across different subjects. We were able to find transitions during which the various networks activate and deactivate correlate with changes in the story which would have been hard to estimate using a model-based approach.

The advantages of the toolkit are multifold. One, as we have discussed, is model-free and does not require a set of assumptions before going into the data except setting the baseline. Second, it not only allows one to analyze block based or event based task design but also naturalistic stimuli based experiments. Third, the toolkit allows for a TR-level analysis of brain activity thus allowing fMRI scientists to find much more interesting spatiotemporal patterns. Fourth, the toolkit allows for flexibility of statistical testing models in addition to region of interest analysis, and contrast analysis that is standard in the field and available in other toolboxes like FSL, SPM, etc., thus making it comfortable enough for cognitive neuroscientists to get accustomed to but providing many insights into their data without a long learning curve.

One limitation of the toolkit is the data requirement. Unlike standard tool boxes which can analyze with one session of data within a subject, the TSA method requires more than eight subjects for inter-subject analysis. A version of the model can also be used for within-subject across-trial analysis which we have called Inter-Trial TSA (IT-TSA) in the past <u>(Tripathi & Garg, 2022)</u> which would again require at least 8-10 trials within each subject. Given the ubiquity of large-scale datasets like HCP, NKIRS, and open sharing practiced widely across the MRI community, we hope that this limitation will be easily overcome. Another challenge that needs to be overcome would be the alignment of brains across individuals. Studies have shown that the transmodal regions that are responsible for higher-order cognition have slight differences across subjects <u>(DiNicola & Buckner, 2021)</u>. It would be important to match networks and regions across subjects using hyperalignment methods to improve the identification of synchronized activity.

We hope that experimental and cognitive neuroscientists would find valuable use of this toolkit which could enable more discoveries in the field of brain mapping and spatiotemporal dynamics of cognition and can allow for more utility of fMRI-based approaches to real-world applications.

## Online Methods

### TSA Toolkit Methods

#### Normalized Signal Change

The Temporal Synchronization Analysis (TSA) measures the synchronization among a set of signals at different time steps. Consider a set of signals, 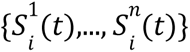 which represent measurements of BOLD signal for ‘*n*’ subjects at a particular voxel ‘*i*’ and time ‘*t*’. For each subject, denoted with *p*, the Normalized Signal Change (NSC) at time *t*, with respect to a baseline signal represented as 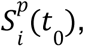 is calculated as follows:

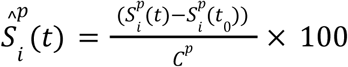

where *C*^*p*^ is a normalization constant obtained by averaging the signals over all voxels and at all times for subject *p*. More variations of the normalization constant and baseline are discussed in the next section. Under the null hypothesis, 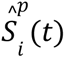 is normally distributed with a zero mean and is independent of all others 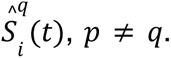 The mean and standard deviation of the NSC, 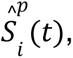 at each voxel *i* and time *t* are estimated as follows:

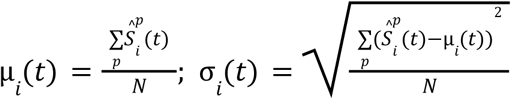

A two-sided t-test is performed for each voxel to obtain the t-statistic that indicates the temporal synchronization of the set of *n* signals at time *t*:

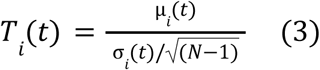

A negative but significant value of temporal synchronization *T_i_* (*t*) at time *t* indicates that the signals for all subjects decrease synchronously at time *t* with respect to their baseline values at time *t*_0_. Similarly, a positive but significant value of *T_i_* (*t*) indicates a synchronous increase. An important parameter in the TSA is the choice of a suitable baseline. In case the experiment contains fixation blocks, the choice of any suitable volume (or the average of such selected volumes) from the fixation blocks that are likely to have no signal bleeding from the previous stimulus may be a good choice.

#### Preparation of Input files

fMRI Data: Registering to a common template, such as the MNI152 template is essential for inter-subject analysis. This can be done using FLIRT, a tool for linear (affine) image registration. Before applying TSA, fMRI data must be registered to a common template; any mismatch in affines between subjects will result in an error. Standard pre-processing steps including motion correction, slice timing correction, temporal filtering, and spatial smoothing need to be performed prior to using the TSA toolkit. The toolkit supports both NiFTI and CiFTI file formats.

Masks: The TSA toolkit allows users to specify a brain mask (0/1 voxel values) in NIfTI or CIFTI format that matches the subject file type. This mask serves two purposes:

- Normalization: If the mask_normalisation option is selected, the specified mask is used to compute normalization (see Baseline and normalization).
- Masking: The mask is superimposed on the output NIfTI files to exclude unnecessary brain areas, such as grey matter and other soft tissues, from the analysis.

#### Baseline and Normalization

The TSA toolkit offers various baseline and normalization options to calculate NSC as defined in Equation 1. Users can specify a list of volume ranges to define the baseline signal. The baseline is computed by averaging the BOLD signal over the specified time ranges. In case Explanatory Variables (EV) files are specified, the baseline is relative to the block onset. In this case, 0 means the volume of task/block onset. -1 means one volume before the task/block onset, -5: -1 means an average of volumes -5 to -1 before task onset. For normalization, the toolkit provides multiple options. These are “voxel” (averages for a single voxel across volumes), “all_voxels” (averages across all voxels across volumes), “mask_normalization” (similar to all_voxels except only the voxels within the mask provided by the user are considered for calculation), “none” (no normalization signal) and “baseline” (normalization signal same as baseline signal).

#### Statistical Testing Models

Before applying any statistical models, the TSA toolkit generates the Normalized Signal Change (NSC) for each subject (Eq. 1) according to the choice of baseline and normalization described above. Subsequently, a T-test is applied to the chosen statistical model to obtain group-level statistics. The toolkit provides three statistical models, each assuming that the NSC of a subject *p*, denoted as 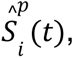 is independent of the NSC of any other subject 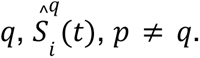

1. Normal - Assumes that, under the null hypothesis, the NSC across all subjects for each time point and voxel follows a normal distribution with a mean of zero. The standard deviation and T-values are as defined in Equations 2 and 3.
2. Binomial - This model assumes that the positive and negative NSC values are equally likely under the null hypothesis. First, NSC values are converted to binary values: +1 if NSC > 0 and -1 if NSC < 0. Two counters are initialized to track the number of positive and negative NSC values (for each voxel and each time) across subjects. The binomial cumulative distribution function (CDF) is then used to calculate p-values for the counter with a higher value.
3. Wilcoxon - The Wilcoxon model in the TSA toolkit tests the hypothesis that the NSC distributions are symmetric around zero using the Wilcoxon signed-rank test, a non-parametric method to test the median NSC across subjects. Once the NSC for all subjects is obtained, 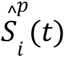 for each from 1 to n, the following steps are used:

– Sort all 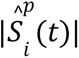 and use this sorted list to assign ranks *R*_1, …,_ *R_n_*: where the smallest observation is ranked one, the next smallest is ranked two, and so on.
– Positive-rank sum *T*^+^ and the negative-rank sum *T*^−^ are used to obtain the Wilcoxon test statistic, *T* as follows:

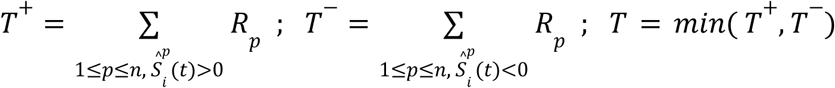
– Calculate the expected mean µ*_i_* (*t*) and standard deviation σ*_i_* (*t*) as follows:

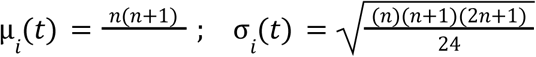

After performing significance testing from one or more of the above statistical models, FDR-corrected p-values (q-values) and uncorrected p-values were saved as output files for each model.

#### Region Of Interest (ROI) Analysis

Atlases/parcellations of the brain can be specified for regions of interest (ROIs) analysis. An atlas file is a NIfTI file where each voxel is assigned a number corresponding to its ROI. If the ROI option is enabled, an atlas file or set of mask files must be provided, and additional ROI-level analysis files are generated as output. For each subject, the NSC is calculated and then processed by the ROI accumulator (see Figure 1). For each ROI in the atlas, the NSC is masked by the corresponding ROI. After masking, the mean NSC of all voxels within that ROI is calculated. This mean NSC for each subject is then used to compute the overall mean and standard error across subjects for each ROI. This ROI analysis can be combined with one of the statistical models described above.

#### EV files, Blocks, Conditions, and their Contrasts

The TSA toolkit allows users to specify EV files, which are text files containing critical timing information about task events during the scan. EV files can be in a 3-column format (used by FSL) or a 4-column format, where the first three columns match FSL’s format (onset time, duration, and value) and the fourth column indicates user-defined condition number for a task event.

- 3-Column EV Files: Each entry is considered a separate block. For example, if a visual stimulus (e.g., a checkerboard) lasts 40 seconds and is presented at t=0 s and t=80 s, it has one condition and two blocks.
- 4-Column EV Files: The fourth column indicates condition numbers. Entries with the same condition number are grouped and assigned incremental block numbers. For example, for two conditions (e.g., pictures of faces and pictures of places), the fourth column would use ‘1’ for face presentations and ‘2’ for place presentations. Each of these conditions can have several blocks.

Contrast between Blocks and between Conditions: The TSA toolkit offers two types of contrasts: Block Contrasts and EV Contrasts. These are optional features that require the EV files to be in a 4-column format and include a weights file specifying the weights of the blocks or conditions. Each row in the weights file corresponds to different contrasts. These contrast signals are generated after the Normalized Signal Change (NSC) has been computed (see Figure 1).

Block Contrast: Block contrast analyzes differences between blocks within the same condition, providing crucial insights when multiple blocks are present. Let the first entry in the weights file be *w*_1_, *w*_2_ ···, *w_b_*, where each *w_j_* is the weight associated with the *j*^*t*ℎ^ block in the EV file (with *b* as total blocks). Let 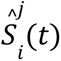 be the NSC calculated for the *j*^*t*ℎ^ block of the EV at the *i*^*t*ℎ^ voxel at time *t* for subject *p*. Note that all these blocks correspond to the same condition, i.e. the entry in the 4^*t*ℎ^ column of the EV files is the same for these blocks. The contrast signal 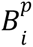 between blocks for a subject p at voxel i is calculated as 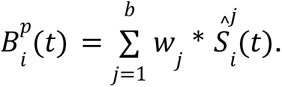 This signal once calculated is passed on to the accumulators for inter-subject analysis as described in the above sections. (See Figure 1)

Condition Contrast: This is the contrast between blocks having different condition numbers but the same block number. Let the first entry in the weights file be *w*_1_, *w*_2_ ···, *w_c_*, where each *w_j_* is the weight associated with the *j*^*t*ℎ^ condition in the EV file (total conditions be c). Let 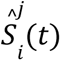 be the NSC calculated for the *j*^*t*ℎ^ condition of the EV at the *i^th^* voxel at time t for subject p. Note all these blocks have the same block number. The contrast signal, 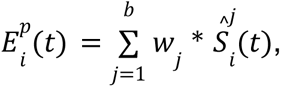 is then passed on to the accumulators for the TSA analysis.

Block Condition Contrasts: The user can apply both these contrasts together as well. In that case, the block contrast files generated would be the input to the Condition Contrast section. Thus, this would result in “Condition contrasts” of the “Block contrasts” which might be useful to analyze in some special cases.

#### Contrast between Groups of Subjects

A second-order analysis could also be run using the toolbox where a user would want to compare activations across groups of subjects. At each time point, a statistical test is run across the distribution of NSC from one group of subjects which can be run using a t-test or other non-parametric tests like Mann-Whitney and Chi-Square test of independence.

### Application of TSA on Datasets

The following publicly available datasets related to different cognitive modalities were used to demonstrate the application of the TSA toolkit:

#### NKI Visual Checkerboard Task Dataset

The NKI-Rockland sample is a large-scale (N>1000), deeply phenotyped dataset for participants across lifespan (ages 6-85 years old) with an aim to accelerate discovery in psychiatry <u>(Nooner et al., 2012)</u>

##### MRI Acquisition

Data were acquired using a Siemens MAGNETOM Tim Trio MRI Scanner. High-resolution MPRAGE T1w images were acquired for the anatomical scans. For the visual checkerboard task fMRI, an EPI sequence was collected with a 32-channel coil (3 mm isotropic voxel size, in-plane FOV = 222 x 222 mm, 40 slices, multi-band factor = 4, TR = 645 ms, TE = 30 ms) with a total of 2.5 mins of data per participant.

##### Pre-processing steps

Standard preprocessing included slice timing correction, motion correction, a high-pass filter (100s), and spatial smoothing (FWHM=5mm). fMRI files were registered to the MNI template using FSL-FLIRT. Subjects missing more than 4% of their brain according to the MNI template mask or with an average RMSE of frame-wise displacement over 1mm were excluded. This resulted in a final count of 317 adult subjects.

##### Experimental Paradigm

The experimental paradigm involved three blocks. Each block commenced with a 20-second fixation period, followed by a task block. During task blocks, participants gazed at a flashing circular checkerboard with alternating white and black squares. The subsequent fixation blocks (15 s) provided no visual stimuli, allowing the HRF to stabilize.

##### TSA Parameters

The TSA for this dataset was conducted with the following parameters.

- Accumulator type - Normal
- Baseline - 0:1
- Normalization - mask_normalization
- Mask - MNI152_T1_2mm_brain_mask
- ROI file/Atlas - Schaefer Cortical Atlas with 400 parcellations and 17 Networks
- EV option enabled with EV files consisting of 3 blocks
- Contrast options are enabled with three options “Block 1 - Block 2”, “Block 1 - Block 3”, and “Block 2 - Block 3”.

### Pieman Narrative Dataset

The Pieman story-listening dataset was acquired as a part of the Narratives collection <u>(Nastase et al., 2021)</u> which aggregated a large number of studies at Princeton University on naturalistic stimuli available publicly on OpenNeuro website (https://openneuro.org/datasets/ds002345/versions/1.1.4).

#### MRI Acquisition

The dataset was collected on a 3T Siemens Magnetom Prisma scanner with a 64-channel head coil. High-resolution T1w scans (1 mm isotropic) were acquired using a single-shot MPRAGE sequence with FoV 176 x 256 x 256 mm. The functional data were acquired using gradient-echo EPI sequence (2.5 mm isotropic voxel size, in-plane FOV = 240 x 240 mm, 48 axial slices, multi-band factor = 3, TR = 1500 ms, TE = 31 ms, total TRs = 300). Further details of acquisition can be found in the dataset publication <u>(Nastase et al., 2021)</u>.

#### Pre-processing steps

The data had a TR of 1.5 seconds. Standard pre-processing steps included slice-timing correction, motion correction, high-pass filtering (cutoff 100s), and spatial smoothing (FWHM=5 mm). Individual fMRI files were registered to the MNI template brain using FLIRT. A dataset of a total of 82 healthy subjects (aged 18-45 years) was used for this analysis.

#### Experimental Paradigm

The “Pieman” audio stimulus (freely available at https://themoth.org/stories/pie-man) is 450 seconds long and begins with 13 seconds of neutral introductory music followed by 2 seconds of silence, such that the story itself starts at 0:15 and ends at 7:17, for a duration 422 seconds, with 13 seconds of silence at the end of the scan. The stimulus started simultaneously with the acquisition of the first functional MRI volume.

#### TSA Parameters

The TSA for this dataset was conducted with the following parameters:

- Accumulator type - Normal
- Baseline - 3:13
- Normalization - mask_normalization
- Mask - MNI152_T1_2mm_brain_mask
- ROI file/Atlas - Schaefer Cortical Atlas with 400 parcellations and 17 Networks

### HCP Working Memory Dataset

The HCP datasets were collected as a part of the WU-Minn Consortium (Van Essen et al., 2013) and comprised 1200 Young Adults collected across multiple sessions including high-resolution structural T1w/T2w scans, resting state fMRI, task-fMRI, diffusion imaging along with a host of cognitive batteries. For this study, we focussed on the working memory task. Description of other tasks and data analysis procedures can be found elsewhere (Barch et al., 2013).

#### MRI Acquisition

High-resolution T1w scans were acquired on a custom Siemens CONNECTOM Skyra MRI Scanner. For the task fMRI, an EPI sequence was collected with a 32-channel coil (2 mm isotropic voxel size, in-plane FOV = 208 x 180 mm, 72 slices, multi-band factor = 8, TR = 720 ms, TE = 33.1 ms, 405 TRs were collected per run) and a total of two runs were collected per participant. Further acquisition details can be found here (Van Essen et al., 2013).

#### Pre-processing steps

for 1200 healthy subjects (aged 18-35 years), already pre-processed files for task data are available on ConnectomeDB (https://db.humanconnectome.org), which follows the minimal pre-processing pipeline in the CIFTI grayordinates space.

#### Experimental Paradigm

consists of 8 task blocks with fixation blocks after every 2 blocks. Within each block are 10 trials of 2 seconds of visual stimulus and 0.5 seconds of fixation. The visual task consists of images of body parts, places, faces, and tools. Subjects are shown the category and some reference images before the onset of each block. They are then asked to identify whether the image is the same as the reference image (0 back) or the image shown earlier (2 back). The stimuli presented are faces, places, and tools. For the TSA results, we used the blocked containing faces as these had a fixation block of 15 sec before the start of the block.

#### TSA Parameters

The TSA for this dataset was conducted with the following parameters:

- Accumulator type - Normal
- Baseline - 0:1
- Normalization - mask_normalization
- Mask - MNI152_T1_2mm_brain_mask
- ROI file/Atlas - Schaefer Cortical Atlas with 400 parcellations and 17 Networks We calculated the 0-back - 2-back contrast between the blocks containing faces.

## Supporting information

Toolbox Documentation

## Declarations

### Ethics approval and consent to participate

Each study in the datasets used in the paper was approved by their respective Institutional Review Boards and all participants were ethically consented.

### Data availability

The HCP Young Adult dataset is available on the platform https://db.humanconnectome.org. NKIRS dataset is available on https://fcon_1000.projects.nitrc.org/indi/pro/nki.html and the Pieman dataset is part of the Narratives dataset release available on the OpenNeuro platform https://openneuro.org/datasets/ds002345/versions/1.1.4

### Code availability

The toolbox is available on Github: https://github.com/Kushagra-Rode27/TSA. And Jupyter notebooks for manuscript figures are available in the same directory.

### Author contribution

SF: Methodology, Formal Analysis, Investigation, Data Curation, Writing - Original Draft, Writing - Review & Editing, Visualization. AD: Methodology, Validation, Formal Analysis, Software, Visualization. KR: Methodology, Validation, Formal Analysis, Software. VT: Conceptualization, Methodology, Supervision, Writing - Original Draft, Writing - Review & Editing. RG: Conceptualization, Methodology, Validation, Writing - Original Draft, Supervision, Funding acquisition.

