## Supplementary Figures for "Temporal Synchronization Analysis: A Model-Free Method for Detecting Robust and Nonlinear Brain Activation in fMRI Data"

### Extended Figures

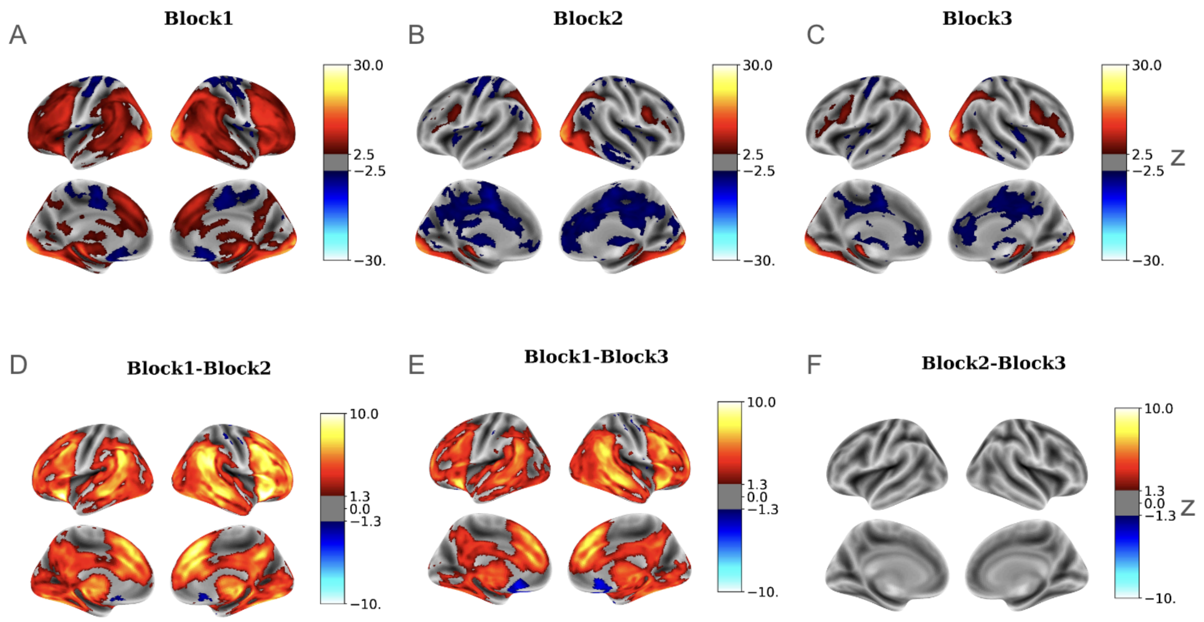


**Fig S1:** GLM analysis results of the visual checkerboard task across 317 participants. FDR-corrected z-statistic maps are shown for (A) Block 1, (B) Block 2, and (C) Block 3. Additionally, FDR-corrected contrast maps are provided for (D) Block 1 vs. Block 2, (E) Block 1 vs. Block 3, and (F) Block 2 vs. Block 3.

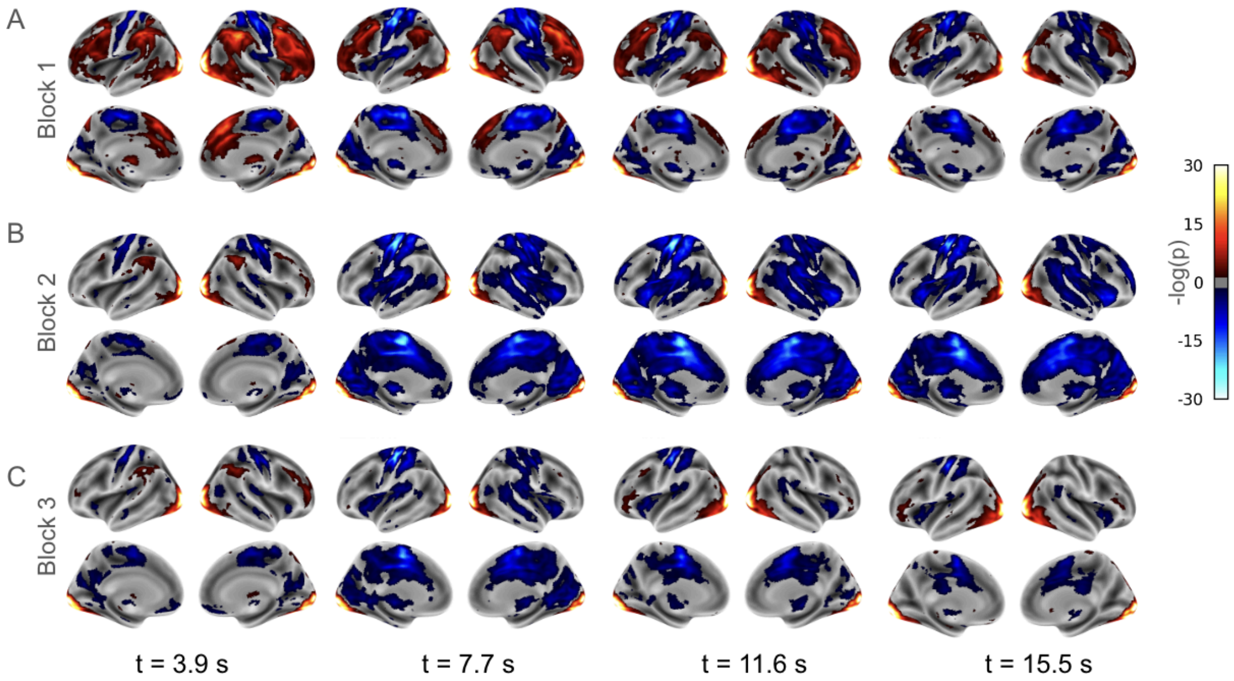


**Fig S2:** TSA of responses to a visual checkerboard task, across 317 adults from the NKI dataset from three different blocks. FDR-corrected p-values (logarithmic scale) for select time points for (A) Block 1 (B) Block 2 and (C) Block 3.

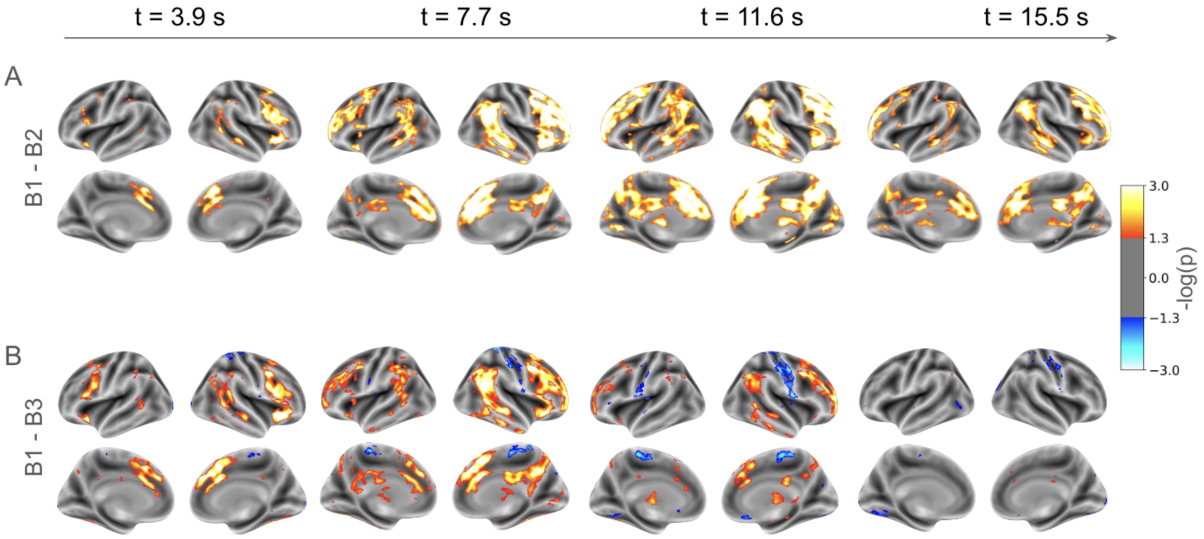


**Fig S3:** TSA results showcasing differences between blocks of the visual checkerboard task across 317 participants. FDR-corrected p-values for select time points for the contrasts (A) "Block 1 - Block 2 (B2-B3)" and (B) "Block 1 - Block 3 (B1-B3)".

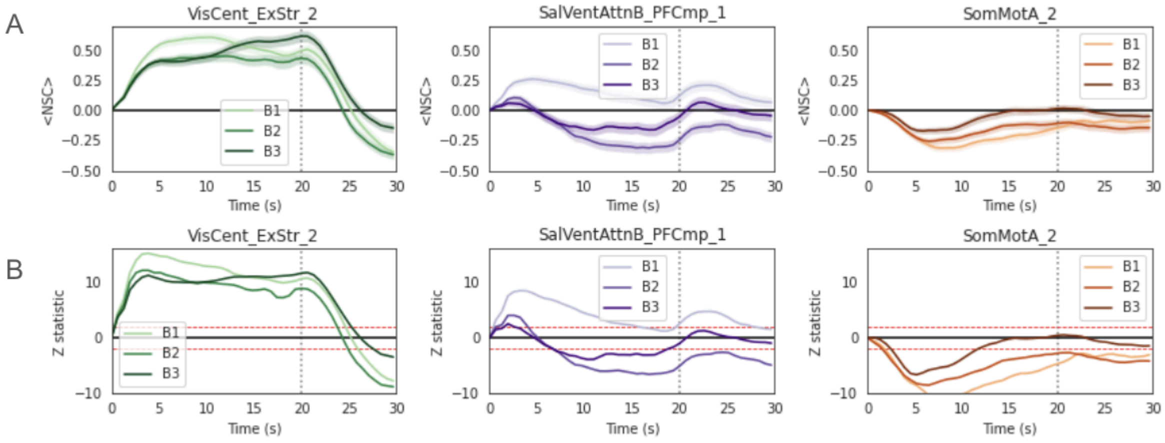


**Fig S4:** ROI analysis of TSA responses to a visual checkerboard task across 317 adults from the NKI dataset, analyzed over three blocks. (A) Normalized signal change over time in three selected ROIs from the Schaefer Atlas. (B) Z-statistics over time in the same three selected ROIs from the Schaefer Atlas. The vertical dotted line marks the end of the visual checkerboard stimulus, indicating the point at which the stimulus ended.

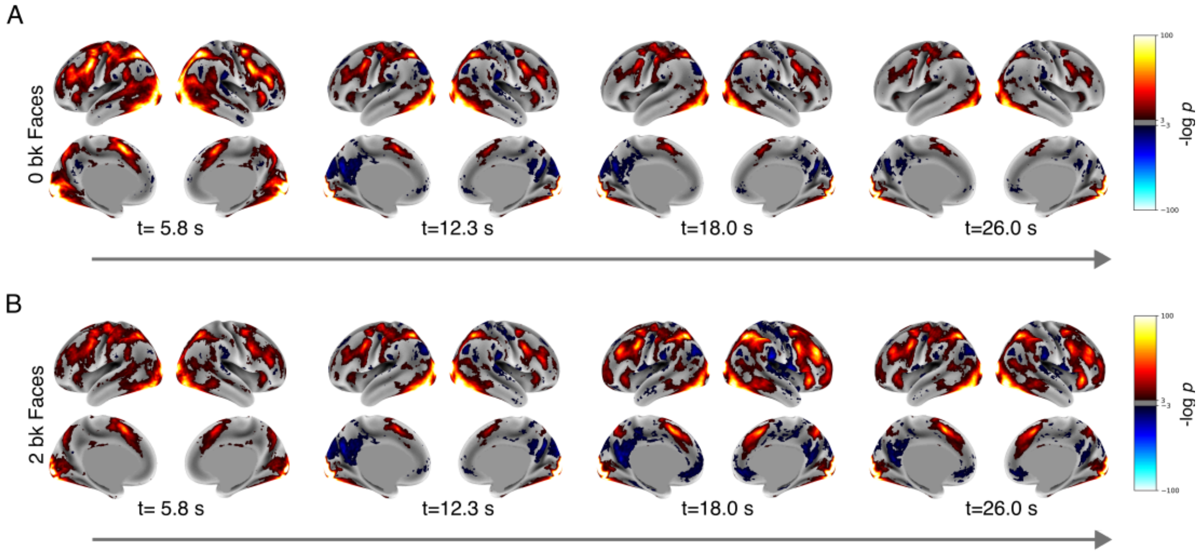


**Fig S5.** TSA results of FDR-corrected p-values for the working memory task (faces as visual stimuli) are shown for (A) the 0-back task and (B) the 2-back task, across selected time points.

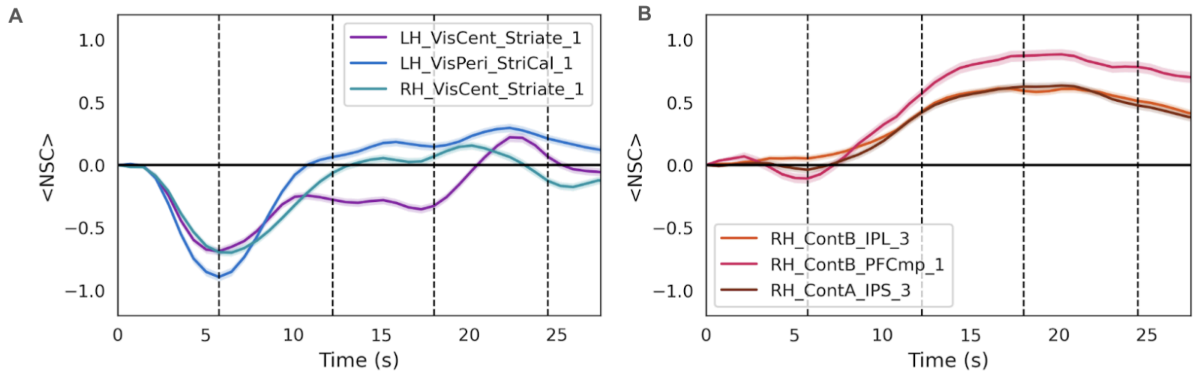


**Fig S6.** ROI analysis of TSA responses for the 2-back vs. 0-back contrasts. (A) Normalized signal change over time and (B) Z-statistics over time for three ROIs with the lowest z-statistics from the Schaefer atlas.

Pie Movie Link :

https://drive.google.com/file/d/1OhSOpmAbfVDesjSzZ4mHDbQgtJlOJyFX/view?usp=sharing
