## Supplementary material for "Temporal Synchronization Analysis: A Model-Free Method for Detecting Robust and Nonlinear Brain Activation in fMRI Data": Toolbox Documentation

### TSA Documentation

#### Structure of a fMRI dataset

fMRI stands for functional Magnetic Resonance Imaging, a special type of brain scan. BOLD stands for Blood Oxygen Level Dependent. It's a type of signal that fMRI scans detect, showing where the brain is more active.

In fMRI studies, there are mainly two types of datasets:

**Resting-state fMRI:**

- What Happens? Here, participants are asked to lie still and try not to think about anything specific.
- Purpose: These scans, taken periodically while the person is resting, help scientists understand how different parts of the brain communicate when a person is not actively doing a task.

**Task-based fMRI:**

- What Happens? In this type, participants are asked to do certain tasks while their brain is being scanned.
- Scanning Process: The scanner takes multiple two-dimensional images at different depth levels (z-positions) of the brain. Then, it combines these images to create a three-dimensional view of the brain.
- Watching Brain Activity: By taking several scans one after the other, scientists can see how the brain reacts to different tasks.

#### Data Formats Explained for Beginners:

**NifTI Format:**

- What is it? NifTI stands for Neuroscience Informatics Technology Initiative. It's a type of file format that replaced an older one called ANALYZE. NifTI is very popular for storing MRI and fMRI brain images.
- What's inside a NifTI file? There are two main parts:
    - Header: This part contains details about the image's size, quality, orientation, and how it fits in space.
    - Data: This is where the actual brain image information is stored.

- Types of NifTI Files:
    - 3D Files: These have three dimensions (x, y, z) representing space.
    - 4D Files: These include time (t) as the fourth dimension, along with the three spatial dimensions.

**CifTI Format:**

- What is it? CifTI stands for Connectivity Informatics Technology Initiative. It was created by the Human Connectome Project to study how different parts of the brain are connected.
- What's special about CifTI? It's lighter and simpler than NifTI. It focuses on the brain's surface, not the whole volume.
- What does a CifTI file contain?
    - Surface Data: It stores information about around 32,000 points on each half of the brain's surface. Each point corresponds to a specific spot on the brain.
    - Volumetric Data: It also includes details about lower parts of the brain like the cerebellum and brain stem.
    - How is it used? You need special software (like ConnectomeWorkbench) to view these maps and understand how different brain areas are linked.

**Metadata and EV Files:**

- What are EV Files? EV stands for Experimental Value. These are simple text files that contain important information about the task being performed during the scan.
- Why are they important? They tell us exactly when each part of the task started and ended for each participant. This timing can vary slightly from person to person.
- Formats of EV Files:
    - 3-Column Format: Includes the time when the task started, how long it lasted, and the EV number (a unique identifier for each task).
    - 4-Column Format: This has an extra column to indicate the specific condition or type of task being performed.

### Task Synchronization Analysis (TSA) of BOLD Signal

- TSA stands for Task Synchronization Analysis. It's a method used to study how signals (like brain signals in an fMRI scan) are synchronized, or how they occur at the same time across subjects.
- The Concept of TSA in Brain Signals:
  - Signals: Imagine you have several signals from different parts of the brain, named S1, S2, ..., Sn. Each signal (like S1, S2) changes over time.
  - Time Points: You look at these signals at a specific moment in time, which we'll call 't'.
  - Baseline (t0): This is a reference time point used for comparison. Think of it as a normal or starting condition for the brain signals.
  - Percent Signal Change: TSA looks at how much the signal's strength has changed in percentage terms compared to the baseline and a normalization constant.
- What TSA Does:
  - Comparing Changes: TSA compares how much each signal has changed from the baseline (t0) to the time point 't'.
  - Two-Sided Test: This is a statistical method to determine if the changes in the signals at time 't' are significant or just random fluctuations.
  - Null Hypothesis: This is a default assumption that there's no significant change. In TSA, the assumption is that the changes in signal are normally distributed (following a typical bell curve) and have an average change of zero.

### Code Usage

usage: `tsa-is [-h] [-f] -i INPUT [INPUT ...] -o OUTPUT [-m MASK] [-d DELTA | -ev EV] [-n] [-b] [-w] [-t TAIL] [-bl BASELINE | -bc BASELINE_CONDITION] [—version] [-ce CONTRAST_EV] [-cb CONTRAST_BLOCK] [-roi ROI_FILE] [-sbsize SUBBLOCKSIZE] [-lm] [-a ALPHA]`

Python-based command-line program for inter-subject temporal synchronization analysis (TSA-IS analysis).

**Required_arguments**:
-i INPUT [INPUT ...], --input INPUT [INPUT ...]
NIfTI input files on which to carry out the TSA-IS analysis
-o OUTPUT, --output OUTPUT
output directory inside which a folder with the name TSA-IS would be created containing the results. It produces 9 files p-value (log and suitable signs), q-value (log and suitable signs), masked q-value(log and suitable signs) t-value (number of subjects that have increased BOLD signal - number of subjects that have decreased for binomial), means, and standard deviations. The remaining two files take the maximum and minimum (across time) of signed log(p) and log(q) values and store them as two volumes: the first volume is the max and the second volume is the min.

**optional arguments**:
-h, --help show this help message and exit

-f, --file If this option is specified, the input is to be given as a file containing the list of files. It is assumed that all the input files are registered to the same atlas.

-m MASK, --mask MASK NIfTI mask file for masking input data. MASK could also be a floating point value describing the threshold, i.e the value of voxels between -threshold to +threshold remains the same and for the rest their values become 0.

-d DELTA, --delta DELTA. Time delta for taking difference (default=1). This option cannot be specified with the -ev option.

-n, --normal consider a normal distribution model on subjects to calculate IISC probabilities (default)

-b, --binomial consider a binomial model on subjects to calculate IISC probabilities

-w, --wilcoxon considers a non-parametric model on subjects to calculate IISC probabilities. The Wilcoxon signed rank test is used for estimating the probabilities.

-t TAIL, --tail TAIL  One-tailed or two-tailed t-test to be performed. TAIL=2 means 2-tailed. TAIL=1 means volume t > volume t - k or baseline. TAIL=-1 means volume t < t - k (or baseline).

-norm NORMALIZATION,  --normalization NORMALIZATION    sets the type of normalization to be used. 5 choices are available - namely, 'none', 'voxel', 'baseline', 'all_voxels', 'mask_normalisation'. NORMALIZATION = 'voxel' averages for each voxel across time. NORMALIZATION = 'all_voxels' averages across time and across the whole brain, i.e averages across all voxels. NORMALIZATION = 'mask_normalisation' is similar to 'all_voxels' normalization, the only difference being that, user specified mask(specified by the -m flag) is applied first over the NIfTI file so as to include only the brain tissues and exclude the other unnecessary parts while computing the normalization. If no mask is specified the standard 2mm MNI_152 mask is used.  NORMALIZATION = 'baseline' makes the normalization values the same as the baseline calculated(for more info, one can take a look at -bl flag and its usage). NORMALIZATION = 'none' as the name suggests indicates no normalization to be used while computing normalized signal change.

-bl BASELINE IF BASELINE is a range, use the mean of the range volumes as a baseline for the tests (-b 3:5,9:20,24:50). This command computes the baseline volume considering the respective time ranges specified and derives the baseline by dividing the summed volumes by the total time points. For instance, if baseline specified as -bl 3:5, 9:20, 24:50, then the total time points would be (5-3) + (20-9) + (50-24) = 39.  In case -ev option is specified, then the baseline is relative to the task onset. In this case, 0 means the volume of task onset. -1 means one volume before the task onset, -5:-1 means average of volumes -5 to -1 before task onset. If no baseline is specified but -ev is specified, the default baseline is -1 (one volume before task onset), and if neither baseline nor ev are specified, the baseline is the average across all volumes. IF BASELINE == 'none', then as the name suggests indicates no baseline to be used while computing normalized signal change. BASELINE == 'none' and NORMALIZATION == 'none' cannot be used simultaneously.

-bc BASELINE_CONDITION  The baseline is the average across all volumes for the given baseline condition number, given in the EV files. EV files must be in the four-column format for this option. Cannot be used with -bl option.

-ev  EV  List of files containing the experimental conditions in the 3- column format of FSL ev files (or 4-column format where the first 3 columns are as in FSL's 3 column EV format and the fourth column is the condition number). In case -f option is specified, then EV is to be interpreted as a file containing a list of ev files for each subject. In case EV files are specified in a 4 column format and no -c (contrast) option is given,  the output is created for each experimental condition and each block separately (also see -o option).

-ce CONTRAST_EV  The CONTRAST_EV is a file with each line representing a different contrast. With this option, the EV files must be in 4 column format. Different entries in a row (line) correspond to the weights assigned to corresponding experimental conditions (first number is the weight assigned to the first experimental condition). So if the first row is -1 1 then the first experimental condition is subtracted from the second experimental condition. The weights assigned in each row need to be in the sorted order of the condition number assigned in the ev files. For example, if the condition numbers are 10, 20 and 30 and the

CONTRAST_EV file contain 1, -1, 0, then contrasts will be taken between condition numbers 10 and 20.

-cb CONTRAST_BLOCK  The CONTRAST_BLOCK is a file with each line representing a different contrast across different blocks of the same experimental condition (or experimental condition contrast). Different entries in a row (line) correspond to the weights assigned to corresponding block (first number is the weight assigned to the first block of the experimental condition). So if the first row is -1 1 then the first block is subtracted from the second block. This option may also be specified  with -ce option. In this case, the output will be "block contrast" of the "EV contrasts". With this option all the block lengths must match, otherwise the results may be meaningless.

-roi ROI_FILE The ROI_FILE is a nifti file of integers containing the region of interest number (roi_number) for each voxel. The roi_number starts from zero to the number of ROIs. This option allows the used to plot the ROI time series (mean, standard deviation) of the brain areas specified by the ROI_FILE. In case -roi option is provided, two csv file containing the mean ROI time series (mean % signal change across the ROI and subjects) and standard deviation of the mean ROI time series (standard deviation across subjects of the ROI-average time series) are also output. In the CSV file, the first column represents the time from the onset of the block and the remaining columns represent the time series of the corresponding ROI number (starting from 1). The voxels belonging to zeroth ROI are ignored.
Make sure the ROI file is properly aligned with the standard MNI mask.

-cr `CONTRAST_ROI` This option has to be given with the -roi option. The CONTRAST_ROI specifies the contrast file (one row per contrast) across the ROIs. The weight of different ROIs are given as different columns in the CONTRAST_ROI files. For example if the roi 4 has to be compared with roi 1 (and the ROI_FILE has six rois) , then the weights in the CONTRAST_ROI file will be -1 0 0 1 0 0 . The output will be csv files (one for each row in CONTRAST_ROI) containing the time series of p-value, t-value, z-value, mean contrast (across subjects and roi voxels), standard deviation of contrast (across subjects, after taking the mean across the voxels of the roi).  The -cr option can be specified with -cr or -cb option to generate roi contrasts of the experimental conditions contrasts or block contrasts.

-sbsize SUBBLOCKSIZE This option is primarily used for running large datasets in limited memory situations. Further, this option is compulsorily used when running the Wilcoxon test due to its large memory requirements. SUBBLOCKSIZE is used to calculate the number of voxels to be allocated to a subchunk of the program. This division of the files into multiple chunks helps with the large memory requirements. The default value is 10.

-lm If specified, the low memory version of normal and binomial tests are run, this flag then requires the user to specify the -sbsize flag. Recommended only when dataset is huge in size. Takes longer time than the usual normal and binomial tests.

-g GROUPS This option has to be given with -cg option. GROUPS should be a .txt file specifying the group name for each subject. Naturally, the number of rows in this file should be the same as the number of subjects. Currently supports 2 different groups. For example -

some subjects can be group A and others can be group B. There must be atleast 2 subjects in each group.

-cg CONTRAST_GROUP This option has to be given with -g option. CONTRAST_GROUP should be a .txt file to specify the configuration for doing group contrast. All the following configurations can be provided together.
"1 1 - disregards groups and saves the TSA output for all subjects "
"1 0 - saves the TSA output only for group 1(lexicographic ordering) "
"0 1 - saves the TSA output only for group 2(lexicographic ordering) "
"1 -1 - runs group contrast (group1 - group2), does not save the TSA output for individual group "
"-1 1 - runs group contrast (group2 - group1), does not save the TSA output for individual group "

-ueqVar This option is to be given with -g and -cg option. Specifies unequal variance 2-sample t-test to be used for doing group contrast. If this flag is not specified then by default the equal variance 2-sample t-test is done for group contrast.

Note - The -g and -cg options are currently available with -n and -w options, -b option does not support these. Further -lm option does not support -g and -cg options currently.

--version show program's version number and exit

This program provides a simple Python-based command-line interface (CLI) for running TSA-IS analysis.

IF no -ev is specified then TSA-IS is computed with respect to the absolute baseline and there is one set of output files for the analysis. In case -ev option is specified, then there is one set of outputs for each experimental block/trial and each condition.

The --input should be two or more 4-dimensional NIfTI (.nii or .nii.gz) files, one for each subject. Alternatively, a wildcard can be used to indicate multiple files (e.g., *.nii.gz).

The --output path should be given as just the directory in which it gives the outputs in the TSA-IS folder in this directory, with prefixes such as ev_<condition_no>_block_<block_no> for different experimental conditions and blocks. In case the EV file is given in 3 column format, then ev_no is always 1. In case the EV file is given in 4 column format and -c (contrast) option is not specified, a separate output file is created for each experimental condition. In case -ce option is specified, then the EV files must be in 4 column format and the output prefix is contrast_ev_<contrast_no>_block_<block_no> and one set of output is generated for every contrast-block pair. In case -cb option is specified, then the output prefix is contrast_block_<contrast_no>_ev_<ev_no>. In case -ce and -cb options both are specified, then the ev file must be in the 4 column format, and the output prefix is contrast_ev_<ev_contrast_no>_contrast_block_<block_contrast_no>. In case -roi option is specified, it outputs two CSV files for each roi (mean and standard deviation), with the prefixes ev_<condition_no>_block_<block_no>

The --mask is again an optional input if masking is to be done.
